# The aftermath of a trophic cascade: Increased anoxia following species invasion of a eutrophic lake

**DOI:** 10.1101/2023.01.27.525925

**Authors:** Robin R. Rohwer, Robert Ladwig, Hilary A. Dugan, Paul C. Hanson, Jake R. Walsh, M. Jake Vander Zanden

## Abstract

Species invasions can disrupt aquatic ecosystems by re-wiring food webs. A trophic cascade triggered by the invasion of the predatory zooplankter spiny water flea (Bythotrephes cederströmii) resulted in increased phytoplankton due to decreased zooplankton grazing. Here, we show that increased phytoplankton biomass led to an increase in lake anoxia. The temporal and spatial extent of anoxia experienced a step change increase coincident with the invasion. Anoxia was driven by phytoplankton biomass and stratification changes, and anoxic factor increased by 10 days. In particular, anoxia established more quickly following spring stratification. A shift in spring phytoplankton phenology encompassed both abundance and community composition. Diatoms (Bacillaryophyta) drove the increase in spring phytoplankton biomass, but not all phytoplankton community members increased, shifting the community composition. We infer that increased phytoplankton biomass increased labile organic matter and drove hypolimnetic oxygen consumption. These results demonstrate how a species invasion can shift lake phenology and biogeochemistry.

**Scientific significance statement:** Invasive species can affect aquatic ecosystems, often by disrupting food webs. We investigated whether the invasive predatory zooplankton spiny water flea could additionally impact the biogeochemistry of a lake, specifically hypolimnetic anoxia dynamics. Using 24 years of observations spanning a spiny water flea invasion that triggered a food web-mediated increase in phytoplankton, we found that increased spring phytoplankton coincided with an earlier onset of anoxia, thereby drawing a connection between a species invasion and a shift in lake oxygen dynamics.

**Data availability statement:** All data is publicly available through the Environmental Data Initiative via identifiers referenced in the methods. Scripts and data to reproduce the results are available on GitHub (https://github.com/robertladwig/spinyAnoxia) and in Rohwer et al. (2023).

**Author contributions:** RRR and RL co-led the entire manuscript effort and contributed equally. RL and RRR came up with the research question and conducted the statistical and numerical analyses: RL analysed the anoxia dynamics and related water quality variables, RRR analysed the phytoplankton community dynamics. RL, RRR, and HAD created figures and visualizations. PCH, JW and JVZ provided essential feedback to the analyses and the discussion of ecosystem implications. RRR and RL co-wrote the paper.

## 1 INTRODUCTION

The introduction of non-native species to lake food webs can disrupt energy flow and mass transfer in aquatic ecosystems, and can threaten aquatic ecosystem stability and services, often to a greater extent than abiotic, anthropogenic environmental changes (Dudgeon et al., 2006; Lopez et al., 2022; Vander Zanden et al., 1999). Many studies of species invasions in lakes focus on food web changes, but the indirect feedbacks species invasions have on lake biogeochemistry are often overlooked. The invasion literature on “zoogeochemistry” is mostly focused on nutrient shunting and relocation. A notable example includes the role of dreissenid mussels in shunting carbon, nitrogen, and phosphorus from pelagic to benthic habitats (Vanni, 2021; Ozersky et al., 2015; Li et al., 2021). However, there are few examples of food web disruptions that lead to alterations in oxygen dynamics in lakes. The paucity of limnological datasets that involve a species invasion and include both lake biology and biogeochemistry has limited our understanding of how species invasions affect foundational biogeochemical processes in lakes.

Lake Mendota is a eutrophic lake in Wisconsin, USA, with a long history of limnological observations through the North Temperate Lakes Long Term Ecological Research program (NTL-LTER). Lake Mendota experienced a population irruption of the non-native predatory zooplankton spiny water flea (*Bythotrephes cederströmii*) in 2009 (Walsh et al., 2016b)^1^. Spiny water flea predation reduced the abundance of the lake’s dominant zooplankton, *Daphnia pulicaria*, which is a keystone species in the food web, and a key food item for native fish populations (Walsh et al., 2016a; Johnson and Kitchell, 1996; Rani et al., 2022). The reduction in *Daphnia* caused a 1 m decline in water clarity due to the reduction in *Daphnia* grazing pressure on phytoplankton (Walsh et al., 2016a), and shortened the duration and intensity of Lake Mendota’s spring clearwater phase (Matsuzaki et al., 2020). The reduction in water clarity was overall due to higher diatom biomass (Walsh et al., 2018), along with an earlier appearance of *Cyanophyta* (Cyanobacteria) during the clearwater phase (Rohwer et al., 2022).

Notable shifts in nutrient concentrations and oxygen dynamics in Lake Mendota have also been observed (Hanson et al., 2020; Ladwig et al., 2022). Ladwig et al. (2021a) applied a mechanistic aquatic ecosystem model that was able to replicate hypolimnetic dissolved oxygen (DO) consumption and bottom-water anoxia from 1995 to 2015, but the model performance declined post-2009, with the model overestimating hypolimnetic oxygen concentrations. This suggests the model did not capture a potential ecosystem shift. Epilimnetic phosphate concentrations from 2010-2018 also decreased by 65% compared to 1995-2009, which could be explained by an increase in whiting events, whereby calcium carbonate precipitation adsorbs available phosphorus (Walsh et al., 2019). In addition, the reduction of grazers (i.e., *Daphnia pulicaria*) and increased phytoplankton biomass may have led to increased phosphorus uptake that was exported to the hypolimnion via sinking algal cells. Together, these studies hint that the irruption of spiny water flea not only reshaped zooplankton and phytoplankton communities, but may also affect seasonal DO dynamics.

Past studies have quantified the impacts of trophic cascades on lake ecosystems (Carpenter and Kitchell, 1993; Carpenter et al., 2001), including on Lake Mendota (Walsh et al., 2016a). Adding to this knowledge, we explore the aftermath of a trophic cascade by quantifying how the impacts of the spiny water flea irruption in Lake Mendota resulted in an increase in the annual spatial and temporal extent of anoxia using 25 years of long-term data. We hypothesize that the spiny water flea irruption caused an abrupt phenological shift in lake anoxia stemming from increased algal biomass. Mechanistically, one could expect increased grazing pressure on planktivorous zooplankton by spiny water flea to cause an increase in spring phytoplankton biomass, which would result in enhanced hypolimentic DO consumption through fallout and mineralisation of algal biomass (Fig. 1).

**FIGURE 1.**
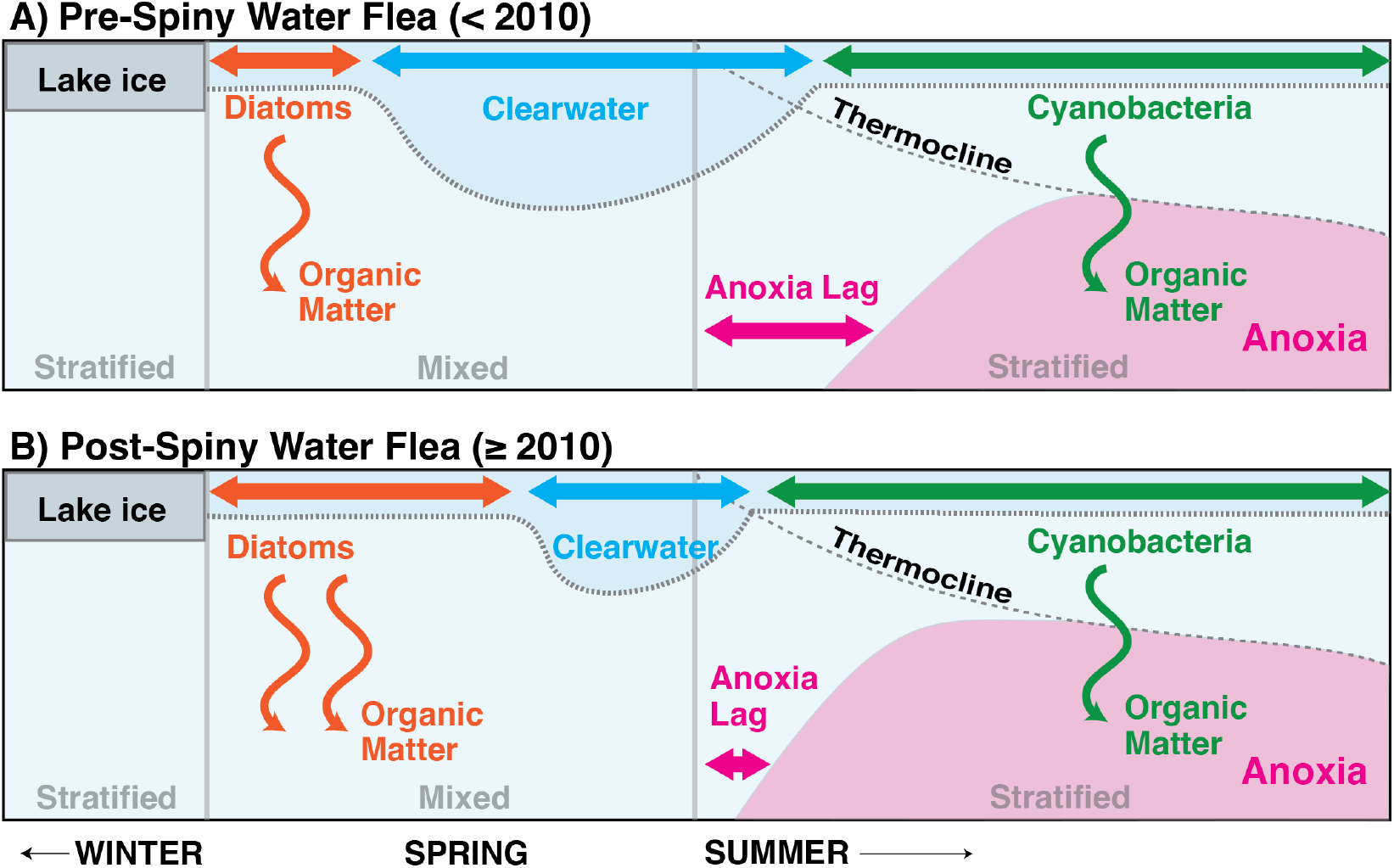
Consequences of the trophic cascade triggered by spiny water flea in Lake Mendota. **(A)Pre-invasion**: Diatom blooms after ice melt are grazed on by zooplankton (esp. *Daphnia*), resulting in a spring clearwater phase that is characterized by deeper Secchi depths (blue shading). After the lake stratifies, hypolimnetic anoxia develops (pink shading) and Cyanobacteria become the dominant phytoplankton. **(B) Post-invasion**: Spiny water flea graze on *Daphnia*, in turn reducing grazing pressure on diatoms. The spring diatom bloom extends and intensifies, and the duration and magnitude of the spring clearwater phase decreases. The additional deposition of organic matter from sinking phytoplankton biomass leads to increased hypolimnetic consumption of oxygen. This reduces the lag-time between stratification onset and the formation of hypolimnetic anoxia.

## 2 METHODS

### 2.1 Lake Mendota

Lake Mendota is a 39.6 ha, dimictic, eutrophic lake with a maximum depth of 25 m (Magnuson et al., 2021a). Physical, chemical, and biological data have been collected fortnightly (when ice-free) to monthly (when ice-covered) by the NTL-LTER since 1995. (Magnuson et al., 2006).

All in-lake measurements were collected in the center of the lake (43.0988N, −89.4054W) and include: ice duration (Magnuson et al., 2021c), integrated water-column measurements from 0-20 m of zooplankton and spiny water flea density measured by zooplankton net tows (Magnuson et al., 2019), integrated water-column measurements from 0-8 m of phytoplankton density and biomass (Magnuson et al., 2022), and depth-discrete measurements of dissolved oxygen (DO), water temperature, nitrate/nitrite 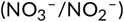, soluble reactive phosphorus (SRP), dissolved reactive silica, and Secchi depth (Magnuson et al., 2021b; Rohwer and McMahon, 2022). Discharge data from the Yahara River were obtained from USGS gage 05427718 (U.S. Geological Survey, 2022).

### 2.2 Data analysis to explore hypolimnetic anoxia

The annual extent of anoxia was quantified using anoxic factor, *AF*, which was calculated according to Nürnberg (1995) as:

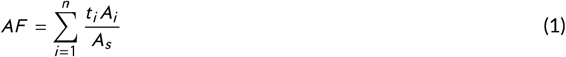

where *t_i_* corresponds to the time duration (days) of an area, *A_i_*, in the water column with DO < 1.5 g m^−3^, and *A_s_* is surface area (m^2^). We chose a conservative threshold for anoxia of 1.5 g ^−3^ (Chapra and Canale, 1991). To identify break points in the time series of annual anoxia, we first applied an Ordinary Least Square Cumulative Sum (OLS-CUSUM) test to quantify the timing of a significant structural change, and afterwards applied the ‘breakpoints’ function from the `strucchange` R-package (Zeileis et al., 2002). Subsequently, years were grouped as either pre- (n = 14) or post (n = 9) irruption, with Jan 2010 as the breakpoint, and Wilcoxon Rank Sum Tests were used to compare groupings.

To evaluate candidate predictors for anoxic factor, we curated datasets of multiple predictors. Biweekly water temperature measurements were temporally interpolated to daily values using linear, constant, and spline interpolation. The transition from mixed to stratified conditions (stratification onset) was defined as when the density gradient between surface and bottom water layers was > 0.1 g m^−3^ and the water column had an average temperature > 4 °C. Stratification duration was quantified as the number of days between stratification onset and offset (when these conditions were no longer met). DO measurements were temporally interpolated using spline interpolation. Acknowledging that these variables do show non-linearity in timeseries, interpolation adds uncertainty to some of the following analyses. However, the inherent autocorrelation in both temperature and oxygen dynamics suggests that over 24 years of data, biweekly interpolated data should result in a high signal to noise ratio. Nutrient data were temporally linearly interpolated to weekly values with the `NTLlakeloads` R-package (Dugan, 2023), area-averaged, and labeled as either surface or deep water based on a mean thermocline depth of 13 m.

Clearwater phase intensity (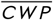 in meter-days) was quantified by integrating Secchi depths between April and June:

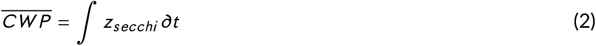

where *z_secchi_* are Secchi depths (m) lineally interpolated to daily values. This method allowed us to quantify year-to-year variability in the intensity of the clearwater phase without the need to define a threshold that would arbitrarily correspond to the formation or breakdown of a clearwater phase.

Interpolated daily DO data were used to calculate the hypolimnetic oxygen consumption fluxes, volumetric, and areal consumption fluxes, according to the Livingstone and Imboden (1996) model. For calculating the vertical anoxia height, only DO concentrations below the bottom depth of the metalimnion, calculated using `rLakeAnalyzer` (see Read et al. 2011) were used.

Candidate predictor selection to explain the inter-annual variability in anoxic factor in a multiple linear model was determined with the ‘Boruta’ random forest classifier function from the `Boruta` R-package (Kursa and Rudnicki, 2010). Candidate predictor importance was calculated using the `relaimpo` R-package *sensu* Lindeman et al. (1980). For the predictor analysis, we included: annual stratification duration, stratification start and break-down date, ice cover dates and duration from the previous winter, summer volumetric, areal, and total oxygen sink, annual days of phytoplankton biomass surpassing concentrations ranging from 0.5 mg L^−1^ to 3 mg L^−1^, annual total Yahara River discharge, annual spring clearwater intensity, maximum Secchi depth during spring, annual average spiny water flea biomass, annual average surface and bottom SRP and nitrate concentrations, and annual average silica concentrations. Important candidate predictors were analyzed using a multiple linear regression model, where a stepwise model selection based on AIC was used to remove predictors.

### 2.3 Phytoplankton and anoxia phenology

Sampling dates were divided annually into four “lake seasons” based on water temperature profiles: 1) ice stratified period, 2) spring mixed period, 3) summer stratified period, and 4) fall mixed period. Phytoplankton biomass within each season and year were averaged to account for uneven sampling and compared between seasons, pre- and post-2009. Changes in phytoplankton community composition were further investigated using the `vegan` R package (Oksanen et al., 2020). Bray-Curtis differences between average annual communities were calculated and analyzed with nonmetric multidimensional scaling (NMDS). Analysis of similarities (ANOSIM) was applied to the distance matrix to determine if years within an invasion group were statistically more similar to themselves than to all years. Shannon and Simpson diversity were calculated for each year and averaged by invasion group to compare pre- and post-invasion diversity. Oxygen phenology was investigated as the difference in days from when stratification developed to when the lowest hypolimnion layer dropped to < 1.5 g m^−3^ DO.

## 3 RESULTS

### 3.1 Anoxia increased with spiny water flea irruption

Anoxic factor increased from an average of 56 days pre-2010 to 66 days, concordant with the spiny water flea irruption in 2009 (Fig. 2A). We explored the temporal trends of potential contributors to anoxia to identify patterns that may be coincident with the 2010 shift in anoxic factor. The phytoplankton-related metrics in general showed significant change (p < 0.01) between pre- and post-spiny water flea regimes (Fig. 2B): modeled average volumetric oxygen consumption increased by 0.03 ± 0.03 g m^−3^ d^−1^ (Fig. 2C, p-value < 0.1 compared to the other phytoplankton-related metrics that are p < 0.01), average total days with phytoplankton biomass >1.0 mg L^−1^ increased by 75.6 ± 20.0 days per year (Fig. 2D), average spring clearwater intensity decreased by 169.1 ± 99.0 meter-days per year (Fig. 2E). Meanwhile, stratification and ice duration did not significantly change(Fig. 2F-G). SRP decreased, with a significant (p < 0.01) decline in the surface layer (Fig. 2H).

**FIGURE 2.**
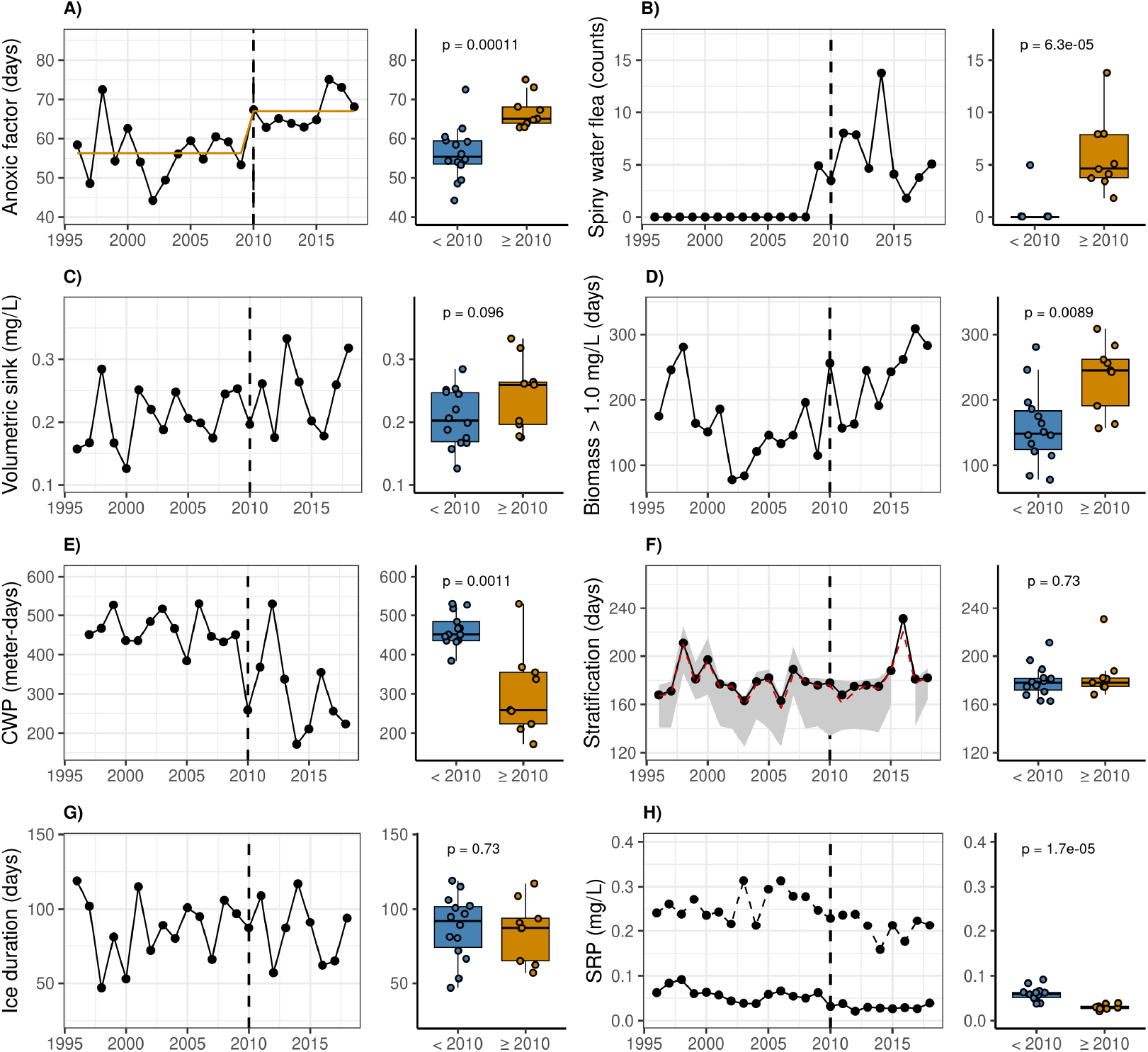
Long-term dynamics of lake variables. **(A)** Anoxic factor over time. Breakpoint analysis of anoxic factor identified 2010 as a breakpoint. The dotted vertical line indicates the breakpoint. **(B)** Spiny water flea abundance over time. **(C)** Modeled hypolimnetic volumetric oxygen depletion flux calculated from observed DO data. **(D)** Days of phytoplankton biomass time series with a biomass over 0.5 mg L^−1^ per year. **(E)** Spring clearwater phase (CWP) over time quantified from Secchi depth. **(F)** Stratification duration over time. Grey ribbon represents the potential uncertainty between sampling points. The red line represents the spline interpolation **(G)** Ice duration over time. **(H)** Annual average SRP concentrations in the surface water layer (solid line) and bottom water layer (dotted line). Box plot highlights only the surface layer SRP concentrations.

Six predictors were determined to be significant predictors for the annual anoxic factor: annual stratification breakdown, annual ice-cover duration, annual duration of phytoplankton biomass over 1.0 mg L^−1^, annual average spiny water flea abundance, annual average surface water layer SRP and bottom water layer SRP concentrations. The resulting linear model has an R^2^ of 0.87 and a p-value < 0.05. Stratification duration, phytoplankton biomass and spiny water flea biomass had a positive correlation with anoxic factor, whereas ice cover duration and both SRP concentrations were inversely correlated. Phytoplankton biomass (38%) and stratification (29%) drove interannual variability in anoxic factor, whereas the remaining four predictors accounted for less than 30% (9% for spiny water flea biomass, 8% for ice duration, 7% for surface SRP, and 5% for bottom SRP).

### 3.2 Spring biomass changes are coincidant with increase in anoxia

After establishing that a step-change increase in anoxic factor occurred in 2010, we further characterized the phenology of anoxia change by investigating the vertical extent of anoxia in the water column before and after the spiny water flea irruption (Fig. 3B). Anoxia expansion started earlier in the year after the spiny water flea irruption, shifting from July to June. However, a shift in the timing of anoxia breakdown was not apparent, nor were changes in the gradient or maximum vertical extent. Given that the linear regression model identified phytoplankton biomass as a driver of the anoxia variability, we compared phytoplankton biomass before and after the spiny water flea irruption (Fig. 3A). Post-invasion phytoplankton biomass was elevated prior to the period of anoxia.

**FIGURE 3.**
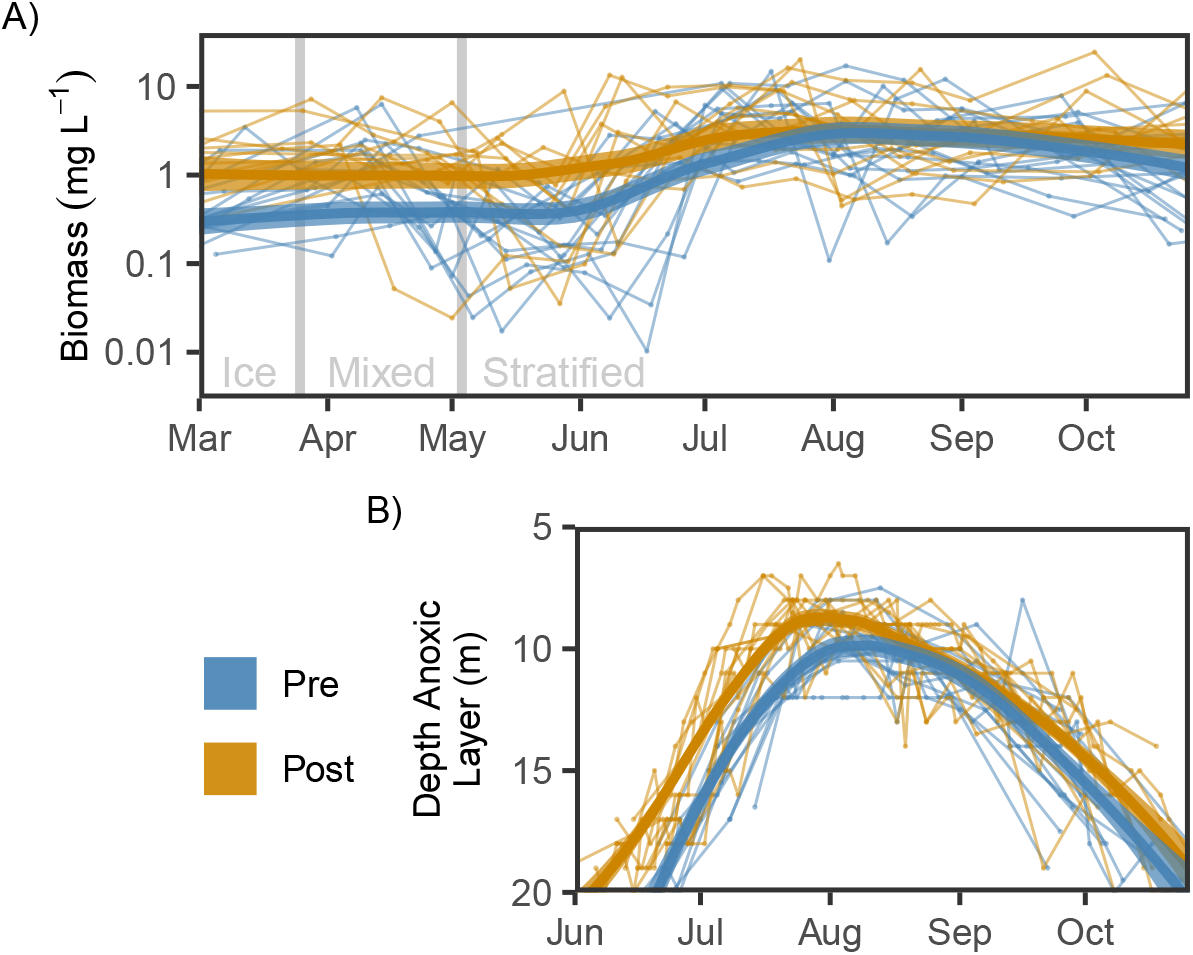
**A)** Annual time series of total phytoplankton biomass before (pre) and after (post) spiny water flea invasion in 2010. Grey lines denote the average timing of ice-off and spring stratification. **B)** Annual time series of anoxia transition depth (DO < 1.5 g m^−3^).

We compared the average phytoplankton biomass within a season for each year before and after the spiny water flea irruption (Fig. 4A). Mean spring phytoplankton biomass increased to concentrations typical of the stratified summer season, from 1 ± 1 (pre-invasion) to 3 ± 2 mg L^−1^ (post invasion) (p < 0.005) (Fig. 4A). Similarly, biomass under lake-ice increased to concentrations previously typical of spring, from 0.3 ± 0.3 to 2 ± 2 mg L^−1^ (p < 0.005). In contrast, later in the season no statistically significant change in total biomass was observed during the stratified summer season (p > 0.1) and more modest increases were observed during the fall mixed season (p < 0.05).

**FIGURE 4.**
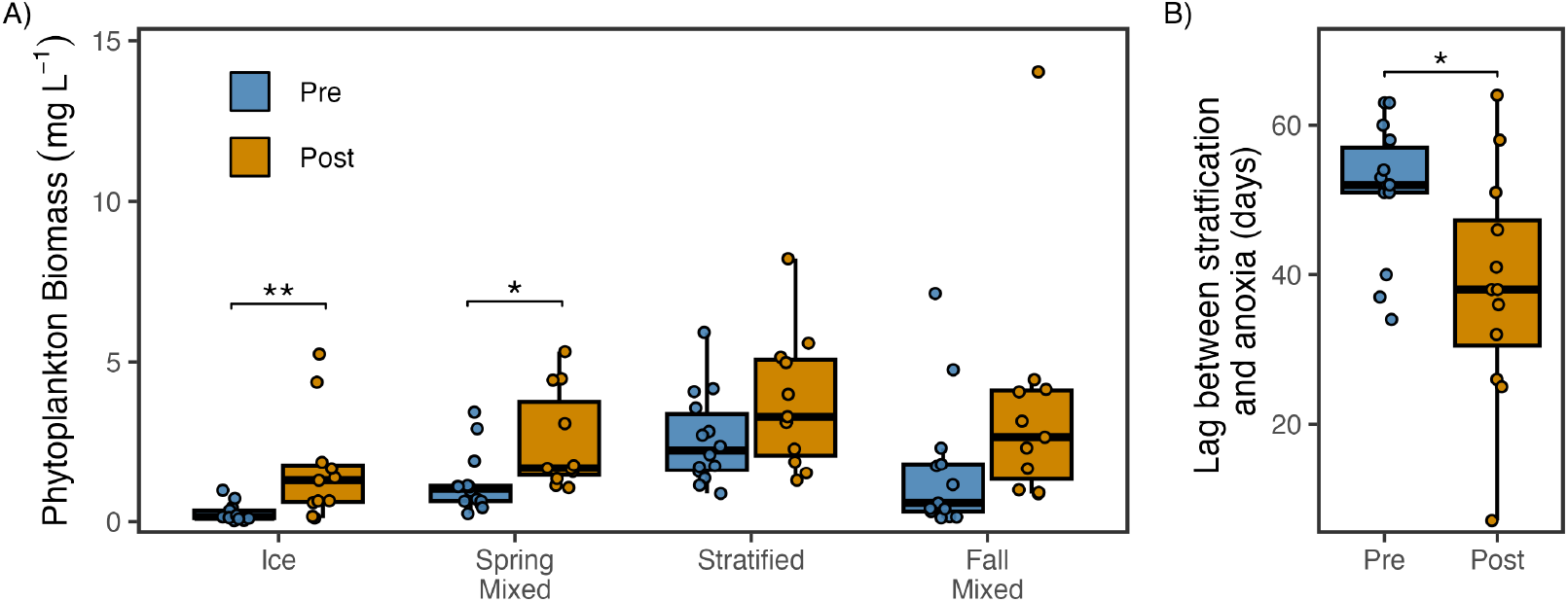
Comparison of seasonal phytoplankton biomass before and after spiny water flea invasion. **(A)** Boxplots of annual averages of phytoplankton biomass in each season. Ice and spring mixed had significantly different phytoplankton biomass post-spiny water flea (p < 0.05, Wilcoxon test with Bonferroni correction). **(B)** Boxplot of lag between stratification development and anoxia onset in days. Anoxia onset occurred sooner after stratification development post-spiny water flea (p < 0.05).

To further examine whether anoxia onset shifted earlier, we calculated the lag between stratification onset and anoxia onset, when DO in the lowest hypolimnion layer was < 1.5 g m^−3^. Post invasion, the lag between stratification and anoxia onset decreased by nearly 2 weeks, from 51 ± 9 days to 39 ± 15 days (p < 0.05) (Fig. 4B).

### 3.3 Diatoms drive phytoplankton increase

Seasonal phytoplankton communities were broadly consistent before and after the spiny water flea invasion; spring was dominated by *Bacillariophyta* (diatoms), summer was dominated by *Cyanophyta* (Cyanobacteria), and fall was dominated by a mix of diatoms and Cyanobacteria (Figure 5). This phytoplankton phenology is typical of a eutrophic lake (e.g., PEG model, Sommer et al. 1986) and previously documented in Lake Mendota (Carey et al., 2016). Diatoms were predominantly responsible for the increase in spring biomass, comprising the majority of the phytoplankton community in all years (67 ± 20% and 65 ± 25% respectively). Diatom biomass in the spring increased 2-fold, from 0.9 ± 0.9 to 2 ± 2 mg L^−1^ (p = 0.08), but the proportion of phytoplankton biomass comprised of diatoms remained relatively constant.

**FIGURE 5.**
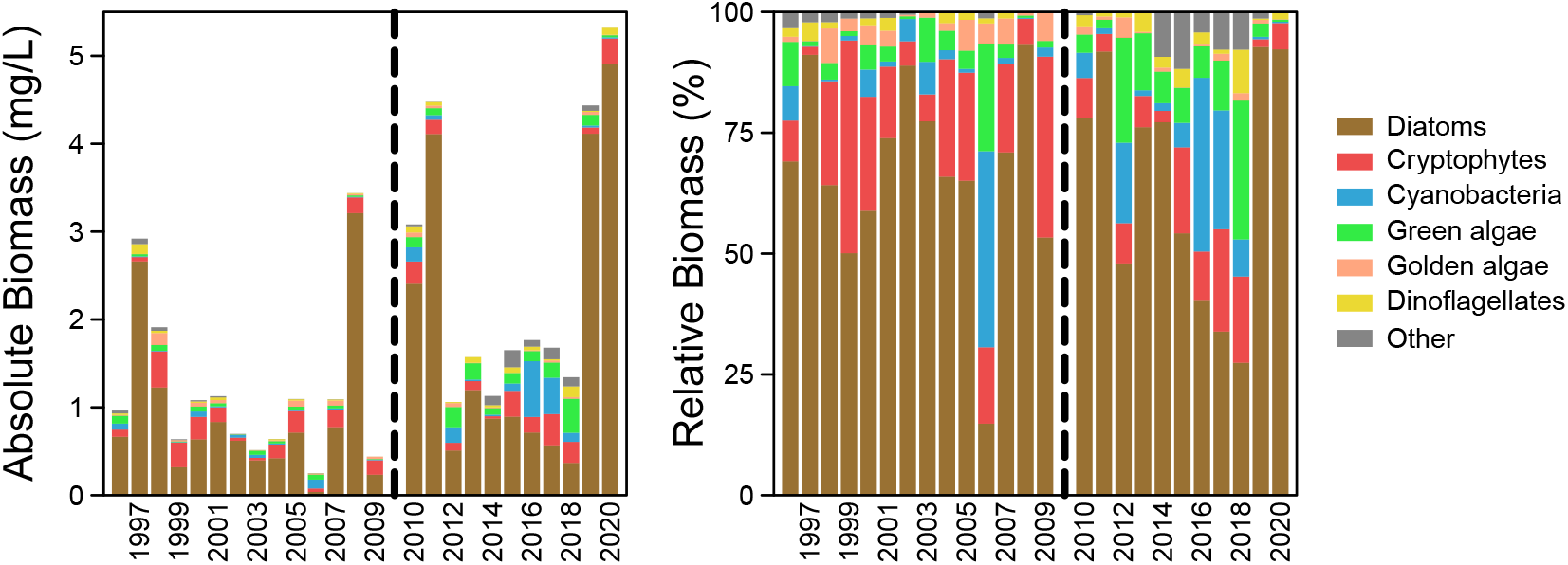
Spring phytoplankton biomass composition. **(A)** Barplots of average annual spring phytoplankton taxa biomass in the spring mixed lake season. **(B)** Barplots of average annual spring phytoplankton taxa relative abundances. The other category includes *Xanthophyta*, *Euglenophyta*, *Haptophyta*, and unclassified organisms. The spring mixed season is majority diatoms (*Bacillariophyta*), which increase along with green algae *(Chlorophyta)*, Cyanobacteria (*Cyanophyta*), and dinoflagellates (*Pyrrhophyta*). Despite shifts in spring phytoplankton community composition to include more green algae, Cyanobacteria, and dinoflagellates, diatoms remained most abundant in spring and drove the increase in spring phytoplankton biomass.

Although diatoms dominated the spring phytoplankton community, three other phytoplankton divisions also contributed to the increase in spring phytoplankton biomass. *Chlorophyta* (green algae) remained at 5-9% of the community but increased 4-fold, from 0.04 ± .02 to 0.1 ± 0.1 mg/L (p < 0.005), Cyanobacteria remained at 5-9% of the community, but increased by 6-fold, from 0.03 ± 0.03 to 0.2 ± 0.2 (p < 0.005), and *Pyrrhophyta* (dinoflagellates) remained at 1-3% of the community but increased 3-fold, from 0.02 ± 0.03 to 0.05 ± 0.03 (p < 0.05). Two phytoplankton divisions decreased their relative contribution, thus shifting the community composition. *Cryptophyta* (cryptophytes) decreased from 17 ± 12 to 9 ± 7% of the community (p = 0.07), and *Chrysophyta* (golden algae) decreased from 3 ± 2 to 1 ± 1% of the community (p = 0.05), although the absolute biomass of both taxa remained constant. A comparison of community composition using Bray-Curtis distance found that the communities were more similar during years with the same invasion status than among all years (ANOSIM significance < 0.05), but these changes were modest enough that phytoplankton Shannon and Simpson diversity did not significantly change.

## 4 DISCUSSION

Contrary to the pre-2009 interannual dynamics of anoxia, which were predominantly driven by changes in water column stability and stratification duration (Ladwig et al., 2021a; Jane et al., 2022), we highlight that the anoxia increase in 2010 was driven by indirect ecosystem impacts of spiny water flea on Lake Mendota’s phytoplankton. Spiny water flea prey on the zooplankton grazer *Daphnia*, and the reduction in *Daphnia* grazing enabled phytoplankton, primarily diatoms, to flourish (Walsh et al., 2017, 2018). The link between epilimnetic phytoplankton biomass and elevated hypolimnetic oxygen consumption has been well established for eutrophic lakes (Paerl, 1988). The increase in springtime phytoplankton biomass observed in this study likely increased the settling flux of organic matter and availability of a labile substrate for hypolimnetic mineralisation. Given that physical factors like stratification did not change following the species invasion, the observed increases in springtime phytoplankton biomass and lake anoxia, as indicated by the decrease in lag between stratification development and anoxia onset, seems beyond coincidence.

Phosphorus dynamics in the lake also shifted following the spiny water flea invasion (Walsh et al., 2019), with a decrease in SRP. While this pattern runs counter to known positive relationships between nutrient availability and phytoplankton biomass (Conley et al., 2009), biophysical processes may provide an explanation. Whiting events can occur when phytoplankton blooms raise eplimnetic pH through the uptake of inorganic carbon, which triggers the precipitation of calcium carbonate and the co-precipitation of SRP (Walsh et al., 2019). Simultaneously, by reducing *Daphnia*, spiny water flea indirectly reduced grazing pressure on phytoplankton, causing spring diatom blooms to persist longer and at higher concentrations. Increased phytoplankton biomass may have also reduced surface layer SRP concentrations due to uptake. This suggests that in addition to increasing hypolimnetic anoxia, the spiny water flea-triggered increase in phytoplankton shifted nutrient dynamics in Lake Mendota.

The cascading impacts of a species invasion highlight the complexity of lake ecosystems, and the far-reaching impacts of a single species addition. These impacts can also extend through time, as a disturbed ecosystem may be more vulnerable to future disturbance (Turner et al., 2020). The susceptibility of Lake Mendota to spiny water flea may stem from a biomanipulation in the 1980s (Walsh et al., 2017), when piscivorous fish were stocked to improve water clarity. One outcome was a food web with low planktivorous fish abundance, a trophic niche which spiny water flea was able to fill. Species invasions can sometimes pave the way for future invasions (Spear et al., 2021) or introduce new synergies with existing species (Simberloff and Von Holle, 1999). In Lake Mendota, zebra mussels invaded in 2015 (Spear et al., 2022), potentially confounding the second half of our post-spiny water flea analysis. In a long-term study of Lake Mille Lacs, a spiny water flea invasion had no net effect on phytoplankton biomass as a simultaneous invasion by zebra mussels compensated for the increased grazing pressure (Rantala et al., 2022). However, in Lake Mendota, Spear et al. (2021) observed no change in water clarity with the zebra mussel invasion. Rohwer et al. (2022) did observe changes in phytoplankton community composition following the zebra mussel invasion, finding an earlier timing of Cyanobacteria onset in the microbial community^2^. However, we did not observe a change in anoxia extent or phytoplankton biomass associated with zebra mussels. Together, shifts in Cyanobacteria phenology (Rohwer et al., 2022) and the shifts in phytoplankton community composition and abundance observed here are examples of ecosystem shifts following species invasions.

In this study, we show how the spiny water flea invasion affected both phenology and biogeochemistry of the Lake Mendota ecosystem. Quantifying the impact of anoxia on Lake Mendota’s ecosystem is a challenge. Anoxic conditions limit the spatial extent of species habitat (Stefan et al., 1996; Magee et al., 2019; Karatayev et al., 2013), can lead to fish kills (Rao et al., 2014), promote biogeochemical redox reactions resulting in sediment release of nutrients and metals (Hupfer and Lewandowski, 2008), and enhance methane emissions (Tranvik et al., 2009). Increases in anoxia represent an additional impact of the spiny water flea invasion that has not been previously accounted for. However, invasions are not the only drivers or stressors that force abrupt change (Turner et al., 2020). Climate change is shortening ice duration (Sharma et al., 2021), increasing water temperature (Woolway et al., 2022), decreasing wind speeds (Magee et al., 2016), and increasing intensity of rain events in the midwestern US (Kucharik et al., 2010); and road salt is shifting lake stratification regimes (Ladwig et al., 2021b). As climate change-caused disturbances increase the stochasticity of aquatic ecosystems, ecological consequences can be far-reaching, from affecting community composition to increasing the extent of oxygen-depleted waters, thereby further restricting organism habitat. Comprehensive long-term monitoring programs that collect observations of food webs, physical characteristics, and biogeochemistry will continue to be essential for studying how these interacting drivers of change impact all aspects of lake ecosystems.

## Acknowledgements

We thank the North Temperate Lakes Long Term Ecological Research program (NTL-LTER) for providing the long-term observational data that made this study possible. We thank Emily Stanley and Mark Gahler for providing up-to-date data and original profile sheets. We thank the 2022 Joint Aquatic Sciences Meeting for providing the coffee break that inspired this manuscript.

## Footnotes

1 Species name changed from *Bythotrephes longimanus* to *Bythotrephes cederströmii*.

2 Note that our lake season “spring mixed” differs from the “spring” season in Rohwer et al. (2022) in that “spring mixed” also includes a large portion of clearwater phase.

